# PP271, a novel bacteriophage primase-polymerase, enables Primer-Free DNA Amplification

**DOI:** 10.64898/2026.11.06.764790

**Authors:** Alejandra Jiménez-Mejías, Susana Guerra-González, Regina Olmos, Itziar Lázaro, Luis Blanco, María Isabel Martínez-Jiménez

## Abstract

Primases are essential enzymes that initiate DNA replication by synthesizing *de novo* primers for replicative DNA polymerases, which ensure accurate genome duplication owing to their high specificity, fidelity, and processivity. PrimPols have emerged as versatile DNA-initiating enzymes that are widely distributed across diverse replication systems, from bacteriophages to humans.

In this work, a novel PrimPol enzyme, PP271, encoded by bacteriophage vB_EfaS-271, was characterized. PP271 belongs to the archaeal–eukaryotic primase (AEP) superfamily and contains the conserved catalytic core, together with a C-terminal PriCT domain. Enzymatic analysis demonstrated that PP271 functions as a *bona fide* PrimPol, i.e. a DNA primase capable of synthetizing short DNA primers. PP271 priming activity is strongly stimulated by *ADP*, which serves as the preferred initiating nucleotide. Additionally, PP271 exhibits relaxed specificity for selecting the priming site, compared to other PrimPols with more restricted template preferences.

TruePrime™ amplification technology combines the *Thermus thermophilus* PrimPol and the highly processive Phi29 DNA polymerase to achieve primer-free DNA amplification. While highly efficient, further improvements in amplification yield, fidelity and product complexity remain possible. Here, we show that DNA primers synthesized by PP271 are efficiently extended by Phi29 DNA polymerase, enabling their coordinated action. Moreover, the short primers generated by PP271 are expected to minimize the contribution of primase-derived errors to the final amplification product while allowing efficient DNA synthesis across a broader range of sequence contexts. Altogether, these findings identify PP271 as a promising alternative for improving primase-dependent DNA amplification.

**Key points:**

- Bacteriophage EFAS271 encodes PP271, a DNA-dependent DNA primase-polymerase
- PP271 preferentially uses *ADP* as the initiating nucleotide during primer synthesis
- PP271 enables primer-free isothermal DNA amplification with Phi29 DNApol

## Introduction

Bacteriophages represent a valuable source of replication enzymes with unique biochemical properties that can expand the molecular biology toolbox. In this context, bacteriophage vB_EfaS-271 infecting *Enterococcus faecalis* clinical strains (Topka-Bielecka et al., 2020), constitutes an attractive model system. This phage belongs to the family *Siphoviridae*, characterized by a non-contractile tail, and displays an efficient lytic cycle, with a short infection period (∼20 min), a latency phase of ∼8 min, and a burst size of approximately 70 phage particles per infected cell (Topka-Bielecka et al., 2020).

The genome of vB_EfaS-271 encodes a distinct set of replication proteins, including a helicase, two endonucleases, a DNA polymerase B-like enzyme, a DnaG-like RNA primase, and a putative DNA primase/polymerase (PrimPol) (Topka-Bielecka et al., 2020). The presence of two different primases suggests an increased flexibility during replication initiation, and a specialized replication machinery potentially adapted for rapid genome duplication and for resistance to replication stress. Specifically, the vB_EfaS-271 genome encodes two primases representing the two major primase superfamilies: a DnaG-like primase belonging to the TOPRIM (topoisomerase–primase) fold superfamily, and a PrimPol-like enzyme from the archaeo-eukaryotic primase (AEP) superfamily. PrimPols are characterized by their dual primase and polymerase activities, enabling *de novo* synthesis of DNA primers using deoxynucleotide triphosphates (dNTPs) (Lipps et al., 2003). Initially identified in archaeal plasmids (Beck & Lipps, 2007), PrimPols are broadly distributed across bacteriophages, bacteria, archaea and eukaryotes, highlighting their evolutionary conservation and functional relevance (Carvalho et al., 2025; García-Gómez et al., 2013; Gupta et al., 2019; Halgasova et al., 2012; Picher et al., 2016; Rudd et al., 2013; Sanchez-Berrondo et al., 2012; Wan et al., 2013; Zhu et al., 2017). In addition to primer synthesis, they have been implicated in mitochondrial DNA maintenance, DNA damage tolerance, DNA repair pathways, and defense against exogenous DNA (Bailey et al., 2019; García-Gómez et al., 2013; Kobayashi et al., 2016; Mourón et al., 2013; Schiavone et al., 2016; Torregrosa-Muñumer et al., 2017).

The catalytic core of PrimPols contains three conserved motifs: motif A, involved in metal ion coordination; motif B, associated with nucleotide binding; and motif C, also contributing to metal coordination (Iyer et al., 2005). These motifs support the binding of two catalytic metal ions (typically Mn²⁺ or Mg²⁺) required for phosphodiester bond formation between the two initiating nucleotides (Calvo et al., 2019) (Velázquez-Ruiz et al., 2023). The priming mechanism involves binding to single-stranded DNA (ssDNA), recognition of initiation sites, formation of a pre-ternary complex with the incoming nucleotide (Boldinova et al., 2021; Martínez-Jiménez et al., 2018), and dinucleotide synthesis, often assisted by auxiliary domains such as Zn-finger (Martínez-Jiménez et al., 2018) or PriCT domains (Iyer et al., 2005).

*Thermus thermophilus* PrimPol (*Tth*PrimPol) in combination with Phi29 DNA polymerase (Phi29 DNApol) has enabled the development of primer-independent DNA amplification systems such as TruePrime™ (Picher et al., 2016). This approach allows whole-genome amplification without synthetic primers and improves coverage compared to random hexamer-based methods (Dean et al., 2001, 2002; Lizardi et al., 1998). However, it presents limitations including sequence bias based in preferred PrimPol initiation sites, large allelic dropout, and inaccurate copy number variant (Deleye et al., 2016; Gonzalez-Pena et al., 2021). In addition, *Tth*PrimPol can synthesize relatively long DNA products (up to 500–800 bp), which may further compromise amplification accuracy.

Here, we characterize a novel PrimPol enzyme from bacteriophage vB_EfaS-271 (hereafter PP271) with properties that may be advantageous for biotechnological applications. This enzyme preferentially uses Mg²⁺ as a cofactor, similar to Phi29 DNA polymerase, and its activity is enhanced by diphosphate nucleotides as initiating substrate. Notably, and in contrast to *Tth*PrimPol, PP271 synthesizes primers of restricted length, a feature that potentially could improve the fidelity of DNA amplification reactions.

## Materials and methods

### Reagents and biochemicals

Inorganic salts, acids, bases and organic compounds were obtained from Merck (Kenilworth, NJ, USA), Sigma-Aldrich (St Louis, MO, USA) and AppliChem (Darmstadt, Germany). Chromatography resins for protein purification included Ni-NTA Superflow (Qiagen, Hilden, Germany) and Heparin Sepharose Fast Flow (Cytiva, Washington, DC, US). Unlabeled ultrapure *NTPs* and dNTPs were supplied by New England Biolabs (Ipswich, MA, USA), while dNTPs used in amplification experiments were obtained from Roche (Basel, Switzerland). Radiolabeled nucleotides *[γ-^32^P]ATP* and [α-^32^P]dCTP were supplied by Perkin Elmer (Waltham, MA, USA). The M13mp18 ssDNA template was obtained from New England Biolabs (Ipswich, MA, USA). DNA oligonucleotides were synthesized by Sigma-Aldrich (St Louis, MO, USA) and IDT (Coralville, AI, USA). Phi29 DNApol was kindly provided by 4basebio (Cambridge, UK).

### PP271 purification

The pET16 vector encoding EFAS-271 PrimPol was synthesized by Genscript (Nanjing, China) and transformed to *Escherichia coli* strain BL21(DE3) cells for protein overexpression. Expression was induced by the addition of 1 mM isopropyl-β-D-1-thiogalactopyranoside (IPTG) to 1 L of log-phase cultures (OD_600 nm_ = 0.5) grown at 37 °C in LB medium supplemented with 100 µg/mL ampicillin. Following induction, cells were incubated at 15 °C for 18-20 hours. Subsequently, cell were harvested by centrifugation (10 min at 8000 rpm, 4 °C) and the cells pellet was weighed and stored at -20 °C. Prior to purification, pellet was thawed and resuspended in 50 mL of buffer A (50 mM Tris-HCl pH 7.5, 700 mM NaCl, 5% glycerol, 5 mM imidazole, 1 mM PMSF, 2 mM β-mercaptoethanol). Cells were lysed by sonication on ice (10 cycles of 20s ON, 40s OFF), and cell debris was removed by centrifugation (30 min, 15000 rpm, 4 °C). The clarified supernatant was loaded onto a Ni-NTA HisPur™ column pre-equilibrated with lysis buffer. The column was washed with 12 volumes of buffer B (50 mM Tris-HCl pH 7.5, 5 % glycerol, 1 mM PMSF, 2 mM β-mercaptoethanol) containing increasing concentrations of imidazole (5, 15, 25, 45 and 65 mM) and decreasing NaCl concentrations (700, 500 and 200 mM). Bound proteins were eluted using buffer C (50 mM Tris-HCl pH 7.5, 5% Glycerol, 1 mM PMSF, 2 mM β-mercaptoethanol, 200 mM imidazole and 200 mM NaCl). Fractions containing PP271 were further purified on a heparin column equilibrated in buffer C. The column was washed with 14 column volumes of buffer D (50 mM Tris-HCl pH 7.5, 5 % glycerol and 200 mM NaCl). Finally, the protein was eluted in buffer containing 1 M NaCl. Fractions containing highly purified PP271 were pooled and adjusted to 50% (v/v) glycerol. Protein concentration was determined by densitometry of Coomassie-stained 15% SDS-PAGE gels using ImageJ (Schneider et al., 2012), with standards of known concentration.

### Site directed mutagenesis

Point mutations at residue Y121 (Y121A and Y121H) were introduced into the pET16 EFAS 271 construct using the QuickChange II Site-Directed Mutagenesis protocol (Kunkel, 1985). The following primers were used: Y121A-sense 5’ CAA ACA ACT GGC AGG GCC GTT GTA GGA TG; Y121A antisense 5’ CAT CCT ACA ACG GCC CTG CCA GTT GTT TG; Y121H sense 5’ CAA ACA ACT GGC AGG CAC GTT GTA GGA TG; Y121H antisense 5’ CAT CCT ACA ACG TGC CTG CCA GTT GTT TG. All constructs were verified by DNA sequencing.

### Sequence alignment

Multiple sequence alignments of PrimPol orthologues were performed using the COBALT (Constraint-Based Multiple Alignment Tool) server from the National Center for Biotechnology Information (NCBI).

### DNA primase assays

DNA primase activity was evaluated using as nucleotide template a circular M13mp18 single-stranded DNA, the indicated oligonucleotide or a total cellular RNA at the concentrations specified in each figure. Reaction mixture (20 µL) contained 50 mM Tris-HCl /pH 7.5), 0.1 mg/mL BSA, 1 mM DTT, 25 mM NaCl, 2.5% (v/v) glycerol, 2 mM MgCl_2_ and PP271 (500 nM). The indicated concentrations of each *NTP* or dNTP were combined with a radiolabelled nucleotide *[γ-^32^P]ATP* or [α-^32^P]dCTP (16 nM, 250 µCi; 3000 Ci/mmol), as specified in each figure. Reactions were incubated at 37 °C for 30 min unless otherwise indicated. Reactions were stopped by the addition of 8 µL buffer (95% formamide loading buffer) and resolved by denaturing electrophoresis on 8 M urea-containing 20% polyacrylamide gel. Reaction products were visualized by autoradiography, and the bands intensities were quantified using ImageJ software.

### DNA polymerase assay

A Cy5 fluorescence-labelled 15-mer oligonucleotide ( 5’GAT CAC AGT GAG TAC 3’) was annealed to a complementary 28-mer template (5’AGA AGT GTA TCC TGT ACT CAC TGT GAT C 3’) by heating at 80 °C for 10 min in a buffer containing 50 mM Tris-HCl (pH 7.5) and 300 mM NaCl, followed by gradual cooling to room temperature. Polymerase reactions were carried out using 20 nM labelled primer–template DNA in a final reaction buffer containing 50 mM Tris-HCl (pH 7.5), 0.1 mg/mL BSA, 1 mM DTT, 25 mM NaCl, and 2.5% (v/v) glycerol and 10 mM MgCl_2_. dNTPs or *NTPs* were added to a final concentration of 100 µM prior to incubation at 30 °C for 20 min with PP271 (200 nM). Reactions were stopped, resolved, and analysed as previously described.

### Primase/polymerase-coupled assay

A Pulse-Chase experiment was performed to analyse the extension by Phi29 DNApol of primers synthesized by PP271 in two consecutive stages (pulse and chase). During the pulse, reaction mixtures (20 µL) contained 50mM Tris-HCl pH 7.5, 50 mM NaCl, 10 mM MgCl_2_, 1mM DTT, 5% (v/v) glycerol and 0.1 mg/mL BSA supplemented with [α-^32^P]dCTP (16 nM, 250 µCi; 3000 Ci/mmol), dATP/dCTP/dTTP (100 µM) and *ADP* 100 µM when indicated. M13mp18 ssDNA (1 ng) was used as temple and incubated at 30 °C during 20 min with PP271 (200 nM). Half of the reaction was then stopped. For the chase phase, a second half of the reaction was supplemented with Phi29 DNApol (50 nM) and the four unlabelled dNTPs (1 mM), to allow primer extension for another 20 min at 30 °C. Then, the samples were processed as described.

### Primer-less multiple displacement amplification of M13mp18 ssDNA

Reaction (20 µL) contained 2 ng/µL of M13mp18 ssDNA, 100 nM of the indicated Phi29 DNApol and 500 nM PP271 in 50 mM Tris (pH 7.5), 5 mM MgCl_2_, 1 mM DTT and 0.1 mg/mL BSA. dNTP were added at 100 µM each and where indicated *ATP* or *ADP* was included at final concentrations of 1, 10 or 100 µM. After incubation at 30 °C during 1 h, DNA amplification (ng/µL) was quantified using the QuantiFluor® dsDNA and ssDNA system and a Quantus^TM^ Fluorometer from Promega (Madison, WI, USA).

## Results

### PP271, encoded by bacteriophage vB-EfaS-271, conserves PrimPol structural features

Looking for alternative PrimPol enzymes with improved biotechnological potential, a BLASTP search using bacterial PrimPol sequence, was performed. This analysis revealed a putative bifunctional DNA primase/polymerase group from virus infecting *Enterococcus faecalis*, among them a 247 amino acid-long polypeptide encoded by the novel *Enterococcus* phage vB_EfaS-271.

To assess whether PP271 retains the characteristic features of PrimPol enzymes, a multiple sequence alignment including bacterial and bacteriophage PrimPols was performed (Fig. 1). The results demonstrate that PP271 conserves the canonical catalytic core of the archaeo-eukaryotic primase (AEP) superfamily, supporting its classification as a PrimPol. In particular, the three hallmark motifs of AEP enzymes—motifs A, B, and C—are clearly preserved. Motifs A and C contain the invariant aspartate residues (D51, D53, and D114 in PP271), required for divalent metal ion coordination at the catalytic site; while motif B includes the conserved histidine residue implicated in binding and positioning the incoming 3′ nucleotide. These conserved residues are essential for catalytic activity and strongly support the *in silico* annotation of PP271 as an active PrimPol. Downstream of the AEP core, PP271 contains a variable linker region, both in length and sequence composition, connecting the catalytic domain to the C-terminal region. The C-terminal segment harbors a PriCT-1 domain, which includes repetitive stretches enriched in glycine, leucine, proline, and glutamic acid residues.

**Fig. 1.**
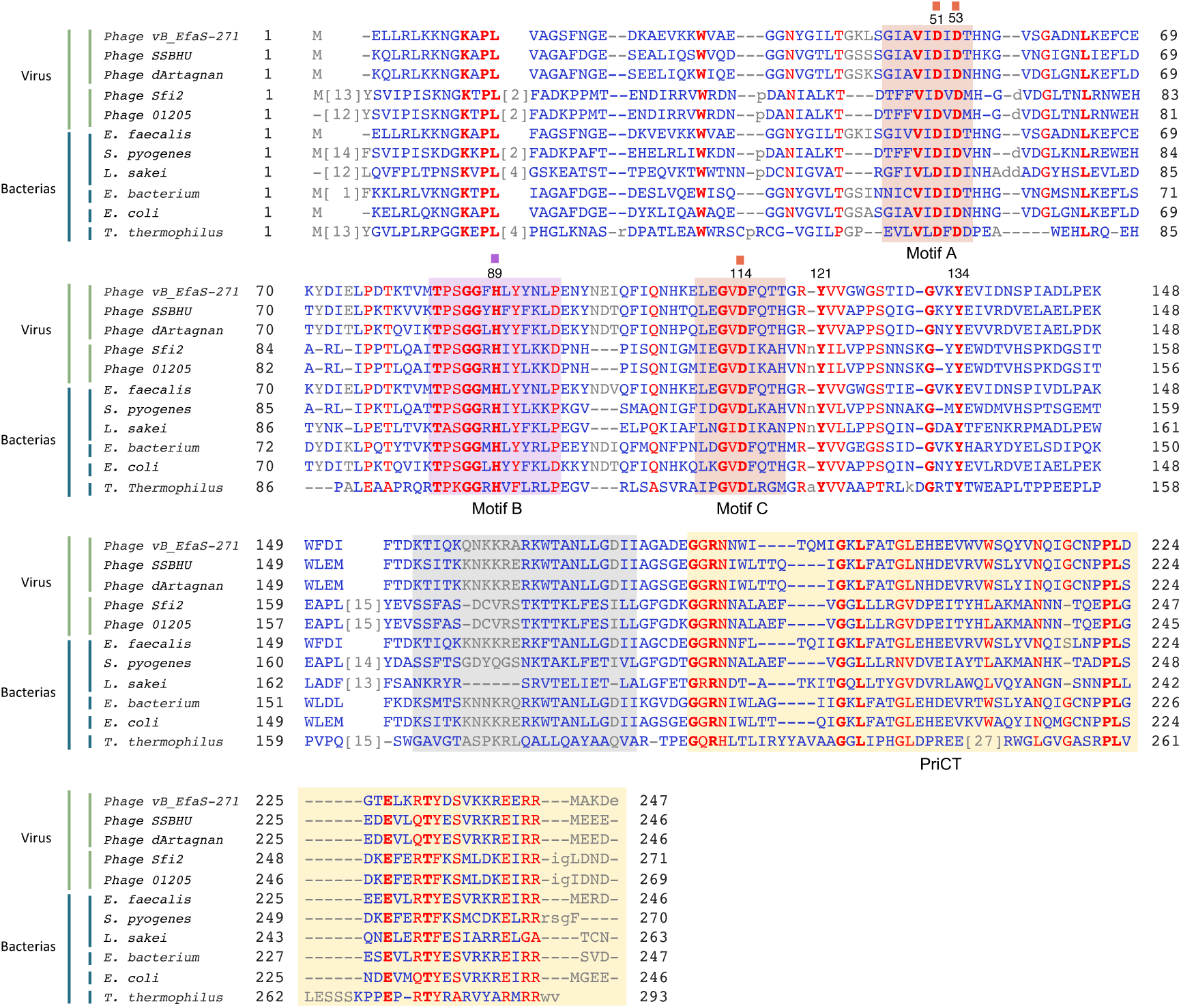
Multiple amino acid sequence alignment of PP271 orthologs. Invariant, partially conserved, and non-conserved residues are shown in red, blue, and grey, respectively. Highly conserved regions characteristic of AEP-like primases are highlighted in red and purple shading. Catalytic residues (in vB_EFaS-271) within the conserved motifs A (D51, D53), B (H89), and C (D11) are indicated by colored boxes above the alignment. The predicted intrinsically disordered linker region is shaded in grey, while the C-terminal PriCT domain is shaded in light yellow. Viral PrimPols: *Enterococcus* phage vB_EFaS-271 (QLF85406.1), *Enterococcus* phage SSBHU (XNX25839.1), *Enterococcus* phage d’Artagnan (CAD7767787.1), *Streptococcus* phage Sfi2 (AAC72437.1), *Streptococcus* phage 01205 (NP_695090.1), bacterial PrimPols, *Enterococcus faecalis* (WP_448890595.1), *Streptococcus pyogenes* (WP_032467158.1), *Latilactobacillus sakei* (WP_112206812.1), *Erysipelotrichaceae bacterium* (MEG2684454.1), *Escherichia coli* (MBL0992195.1), *Thermus thermoplilus* (WP_011173100.1).

This structural and functional conservation, together with its phage origin context, highlights PP271 as a promising candidate for biotechnological applications, including primer-independent DNA amplification strategies and whole-genome amplification workflows.

### PP271 synthesizes DNA primers

PP271 was cloned, expressed, and purified in a soluble and active form, as described in the Methods section. Its primase activity was first evaluated using a single-stranded circular M13 DNA template (Scheme Fig. 2a) in the presence of *NTPs* only. As shown in Fig. 2b, no *RNA* primer synthesis was detected under these conditions, indicating that PP271 does not function as a canonical *RNA* primase. Notably, the vB_EfaS-271 phage genome encodes a DnaG-like primase, suggesting that *RNA* primer synthesis is likely carried out by this enzyme, while PP271 may perform a distinct and complementary function.

**Fig. 2.**
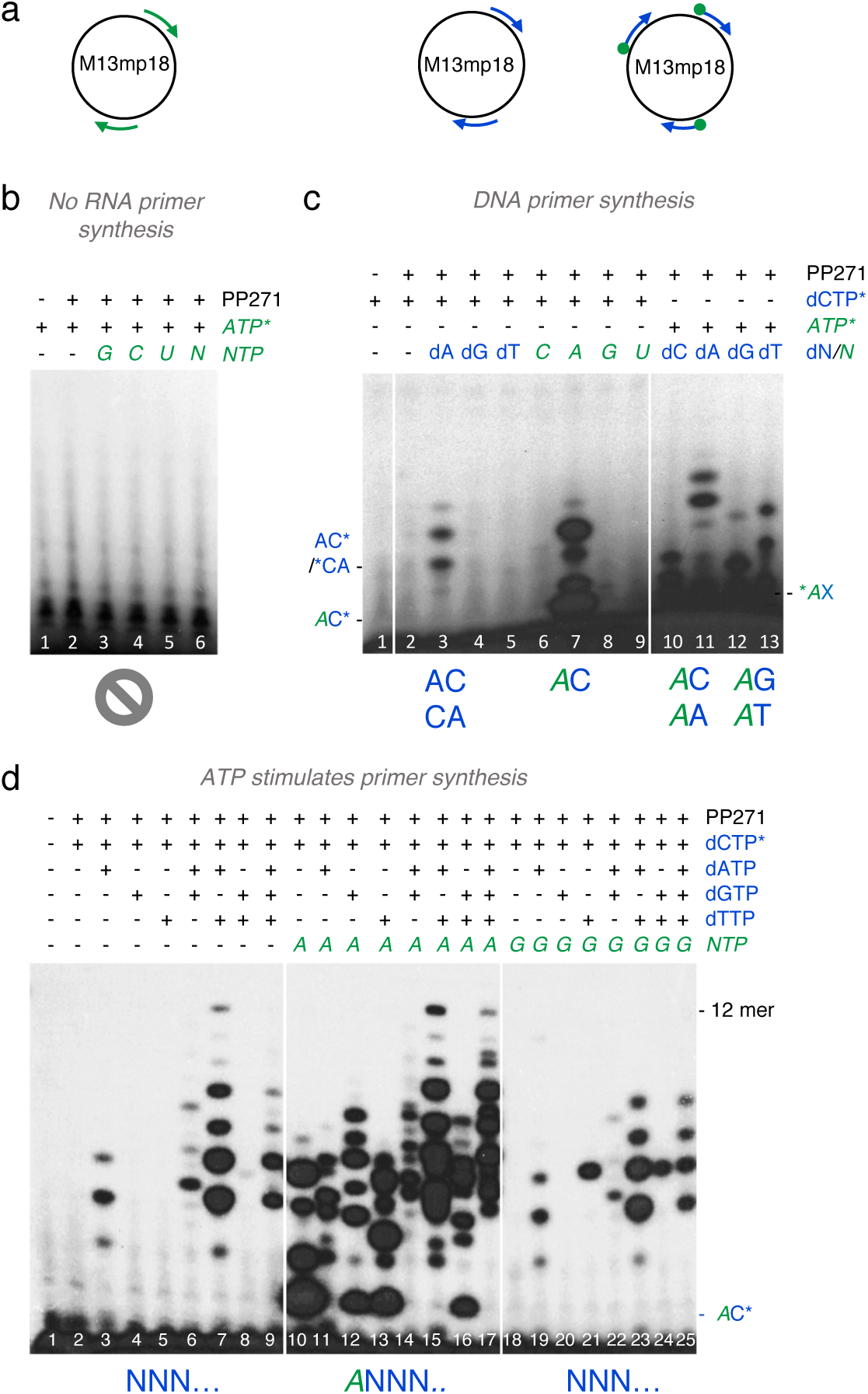
*ATP* stimulates DNA primase activity of PP271. (a) Schematic representation of the possible reactions catalyzed by PP271: primer synthesis on a single-stranded DNA template either with *NTPs* (green arrows), dNTPs (blue arrows) or with *ATP* as the initiating nucleotide followed by elongation with dNTPs (green dot and blue arrow). (b) PP271 (500 µM) does not exhibit *RNA* primase activity, as no *RNA* primer synthesis is detected when only *NTPs* are supplied. (c) Left, PP271(500 nM) synthetizes primers using [α-³²P]dCTP (16 nM) combined with individual dNTPs or *NTPs* (100 µM), showing higher activity when *NTPs* are included. Right, PP271 (500 nM) initiates primer synthesis with *[γ-³²P]ATP* (16 nM) in combination with any of the four dNTPs (100 µM) demonstrating the use of *ATP* as the initiating nucleotide. (d) Left, sequential primer synthesis products generated by PP271 in the presence of dNTPs (100 µM). Center, primer synthesis by PP271 (500 nM) is enhanced in the presence of *ATP* (100 µM), indicating a stimulatory effect when using *ATP* as initiating nucleotide. Right, no effect on primer synthesis by PP271 by addition of *GTP*.

PrimPol enzymes are characterized by their ability to use dNTPs for primer synthesis at both the initiation and elongation sites. However, some members, such as human PrimPol, display a preference for incorporating a ribonucleotide at the 5′ site. To determine whether PP271 is a strict DNA primase using exclusively dNTPs or whether it incorporates a ribonucleotide at the initiation site, dimer formation assays were performed (Fig. 2c). All possible combination of dNTPs or *NTPs* were tested in the presence of one radiolabeled nucleotide ([α-³²P]dCTP or [*γ-³²P]ATP*). PP271 efficiently synthesized primers using dNTPs, confirming its classification as a PrimPol. In addition, PP271 displayed a strong preference for adenine nucleotides, particularly for *ATP* at the 5′ position, as this condition yielded the highest levels of dimer formation. The most favorable nucleotide combination corresponded to the formation of *_3p_A*dC dimers. However, differences in yield were detected depending on the labeling strategy ([α-³²P]dCTP *vs [γ-³²P]ATP*), with higher levels detected when *ATP* was present at µmolar concentration (Fig. 2c, lanes 7 and 10). These results suggest that *ATP* availability may be a limiting factor for dimer formation.

Based on these initial preferences, primer synthesis was further analyzed using dCTP as the labeled nucleotide in combination with the remaining dNTPs, with or without *ATP* as the 5′ initiating nucleotide and using *GTP* as control. As shown in Fig. 2d, *ATP*, but not *GTP*, increases both the abundance and diversity of primers depending on the available dNTPs. Notably, this increment in diversity does not affect primer length, as primers display a similar size distribution under both conditions (approximately 10 nt).

### PP271 DNA primer synthesis is stimulated by *ADP*

Generally, the AEP catalytic core alone is sufficient to support DNA polymerase activity; however, priming requires an additional domain to stabilize binding of the initiating *5′ NTP*. In human PrimPol, this function is performed by a C-terminal Zn-finger domain (Keen et al., 2014; Mourón et al., 2013), whereas bacterial PrimPol contains an orthologous PriCT domain in the same region (Picher et al., 2016). The Zn-finger domain mediates in the interaction with the initiating 5′ nucleotide and promotes the preferential use of a triphosphorylated ribonucleotide (Martínez-Jiménez et al., 2018) in a similar manner as p58 from the human Polα-primase (Baranovskiy et al., 2016). The triphosphate group of the initiating nucleotide not only favors efficient dinucleotide synthesis but also contributes to proper primer elongation (Martínez-Jiménez et al., 2018).

To evaluate the contribution of the phosphate groups of the 5′-site nucleotide in dimer formation by PP271, adenosine nucleotides containing one, two, or three phosphates, in both deoxy-(dAMP, dADP, dATP) and ribo-(*AMP, ADP, ATP*) forms, were incubated with [α-³²P]dCTP which binds at the 3’site (Fig. 3a). As shown in Fig. 3b, no dimer formation was detected with monophosphate adenosine species (lanes 2 and 5). In contrast, diphosphate forms of adenosine, both ribo and deoxy (*ADP* and dADP), promoted the highest levels of dimer formation with [α-³²P]dCTP (*_2p_A*–dC* and _2p_dA–dC*) (Fig. 3b, lanes 3 and 6). *ATP* also promoted not only dimer formation but also elongation to a pentamer probably by the use of *ATP* at the elongation site.

**Fig. 3.**
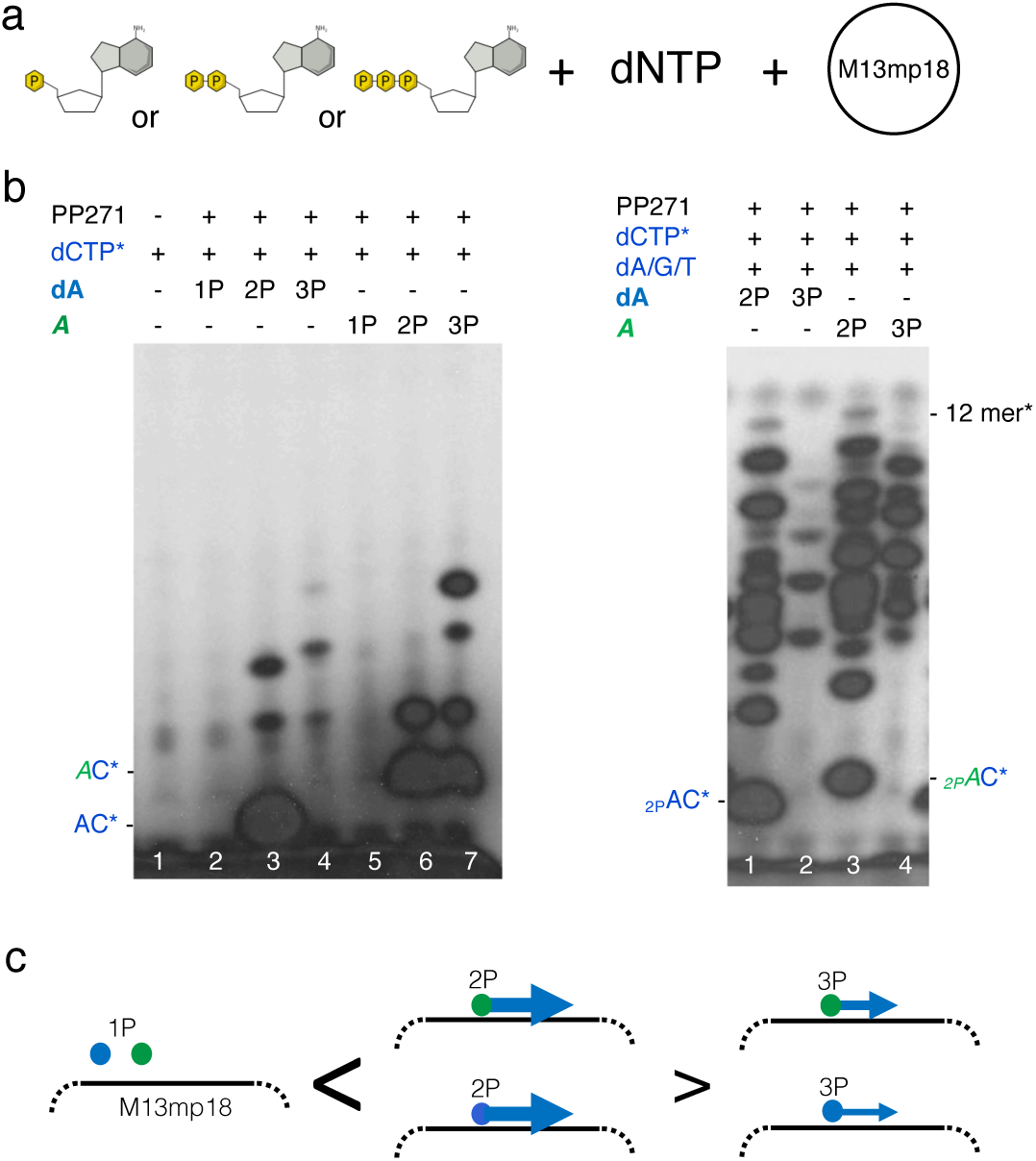
Diphosphate nucleotides are preferred initiators and stimulate primer synthesis by PP271. (a) Schematic representation of the primer synthesis reaction catalyzed by PP271 on M13mp18 in the presence of dNTPs supplemented with mono, di and triphosphate nucleotides. (b) Left, dimer synthesis by PP271 (500 nM) using [α-³²P]dCTP (16 nM) and dA or *rA* with one, two or three phosphates (100 µM). Right, primer synthesis by PP271 using [α-³²P]dCTP (16 nM) together with dATP, dGTP and dTTP (100 µM) in the presence of either dADP, *ADP* or *ATP* (100 µM), as indicated. (c) Schematics showing the efficiency of primer generation (by arrow size), the initiating nucleotide, depicted with a blue (deoxynucleotide) or green (ribonucleotide) circle and 2P or 3P to indicate the number of the nucleotide phosphates.

To determine whether these dimers initiated with dADP, *ADP* or *ATP* could be further extended by PP271, the remaining deoxynucleotides (dA, dG, or dT) were added. *ATP* (Figure 3B, lane 4) promoted primer elongation; however, dADP and *ADP* proved to be the most effective initiator, supporting the synthesis of primers up to 12 nt in length (Fig. 3b, lane 1 and 3). Taken together, these results indicate that *ADP* and dADP are the preferred initiating nucleotide for PP271, as they not only promote efficient dinucleotide formation but also enables subsequent primer elongation.

### PP271 DNA primer synthesis is DNA-templated

So far, PP271 has been shown to function as a DNA-dependent DNA primase. However, a DNA primase able to use *RNA* as template would be of significant biotechnological interest. Therefore, we evaluated whether PP271 is capable of using *RNA* as a template.

As shown in Fig. 4a (lanes 2 and 3), primer synthesis by PP271 on M13mp18 ssDNA was stimulated by *ADP*, consistent with previous observations. In contrast, total RNA extracted from cells did not support primer synthesis, even in the presence of *ADP* (lanes 4 and 5). To further assess template specificity, a 60-mer oligonucleotide (previously used with human PrimPol) was employed. This DNA template contains a central heteropolymeric sequence flanked by poly(T) tails (T_20_GTCAGACAGCAT_29_), along with its *RNA* counterpart (*U*_20_*GUCAGACAGCAU*_29_) as a control. Primer products of approximately 6–8 nt were detected only with the DNA template, and synthesis was stimulated by *ADP*. However, the *RNA* version of the template failed to support priming. These results indicate that PP271 is unable to use *RNA* as a template and therefore lacks reverse transcriptase activity.

**Fig. 4.**
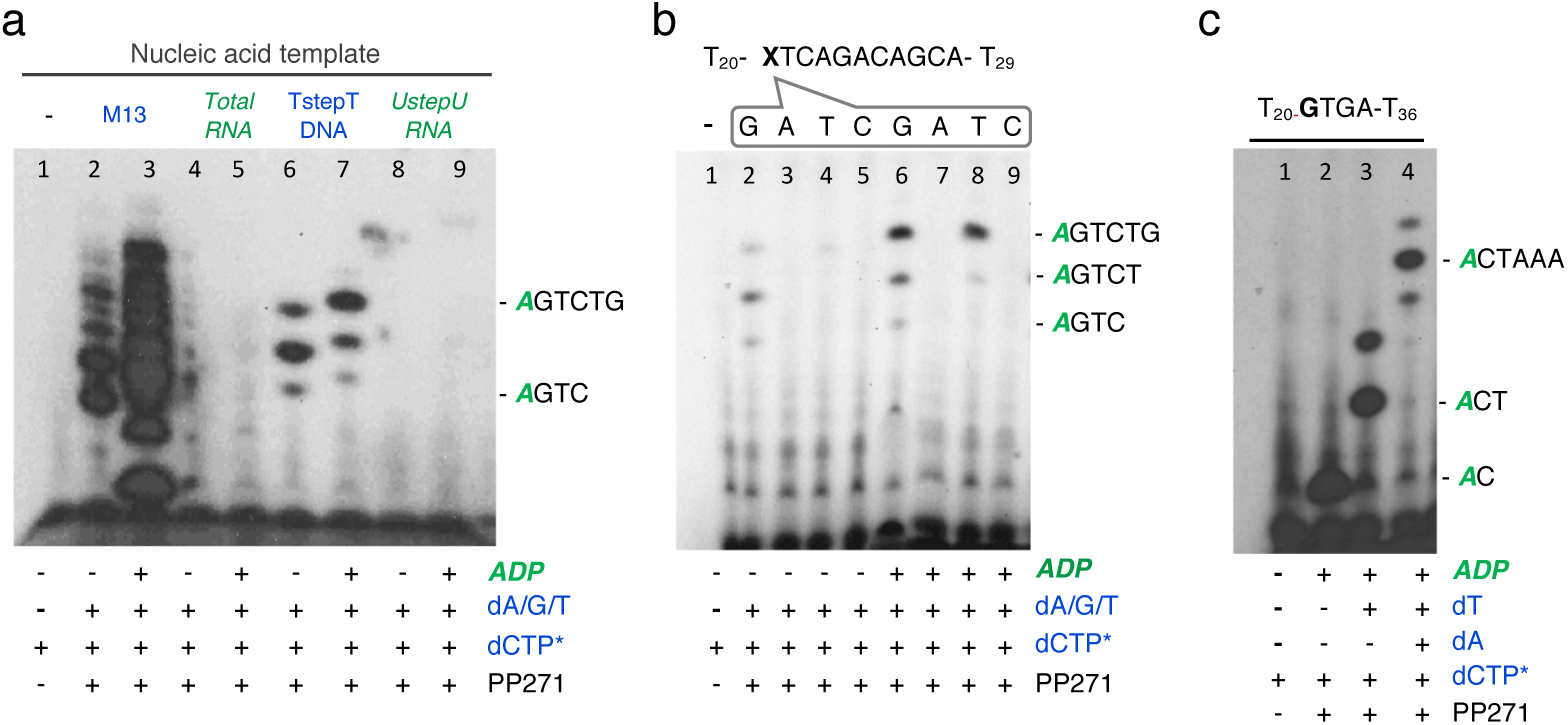
PP271 is a DNA template-dependent DNA primase. (a) Primer synthesis by PP271 (500 nM) using different nucleic acid templates (1 µM): M13mp18 ssDNA (positive control); total cellular *RNA*; DNA oligonucleotide TstepT; *RNA* oligonucleotide *UstepU*. Reactions contained dATP, dGTP and dTTP (100 µM each) and [α-³²P]dCTP (16 nM) and 2 mM MgCl_2_ as metal cofactor. (b) Analysis of the cryptic base preference of PP271 using the defined DNA template TstepT, where X represents any of the four nucleotides. Reaction were performed as described for panel (a) with indicated nucleotide combinations. (c) PP271 incorporates nucleotides during primer synthesis directed by the DNA template sequence. Reactions were performed as described for panel (a) with the indicated nucleotide combinations.

This substrate was used also to analyze template sequence preference during priming. In human PrimPol, like in other primases, initiation involves recognition of a preferred priming site, including the “cryptic base,” defined as the nucleotide preceding the first templating base (Frick & Richardson, 2001). Human PrimPol shows a preference for guanine at this position (Martínez-Jiménez et al., 2018), whereas *Tth*PrimPol prefers cytosine (Picher et al., 2016). To determine whether PP271 exhibits a similar preference, a series of 60-mer oligonucleotides (3′-T₂₀– **X<u>TC</u>**AGACAGCA–T₂₉-5′) containing a central motif (3′ **X<u>TC</u>** 5′), where X represents any nucleotide, was used (Fig. 4b). PP271 showed a preference for guanine (lanes 2 and 6) or thymine (lanes 4 and 8) as the cryptic base, and in both cases, primer synthesis was enhanced in the presence of *ADP*.

To confirm that primer synthesis is template-directed, an oligonucleotide with a unique initiating sequence slightly different to the previous one (T_20_**G<u>TG</u>**AT_36_) was used, allowing stepwise monitoring of elongation from the initial dimer using labeled dCTP. As shown in Fig. 4c, primer length increased progressively upon sequential addition of the required dNTPs, demonstrating that primer synthesis by PP271 is guided by the template sequence.

### Tyr^121^ stabilizes nucleotide binding during PP271 elongation

PrimPols are AEPs capable of synthesizing DNA primers, in contrast to conventional *RNA* primases. In human PrimPol, Tyr^100^ functions as the nucleotide sugar selector, thereby conferring DNA primase activity. Notably, the cancer-associated Y100H variant relaxes sugar selectivity, allowing incorporation of both ribo- and deoxyribonucleotides and enabling the enzyme to act as both a DNA and *RNA* primase. This dual activity has been associated with a selective advantage under replication stress conditions, such as hydroxyurea treatment (Díaz-Talavera et al., 2019) Sequence alignment (Fig. 1) identified two conserved residues in PP271—Tyr^121^ and Tyr^134^— located immediately downstream of motif C, as potential candidates for a similar role in sugar selection. Structural analysis based on the AlphaFold model of PP271 supported Tyr^121^ as the most likely residue to interact with the sugar moiety of the incoming 3′ nucleotide. This spatial arrangement is consistent with a role in nucleotide binding and sugar discrimination, potentially contributing to the control of *NTP versus* dNTP incorporation.

To confirm this hypothesis, two mutants were generated: Y121A, expected to abolish side-chain interactions with the incoming nucleotide, and Y121H, designed to remove the hydroxyl group while partially retaining aromatic character and mimicking the human PrimPol Y100H variant.

Primase activity was first evaluated using an M13mp18 ssDNA substrate in the presence of either dNTPs or *NTPs*, with primer synthesis monitored by incorporation of labeled dCTP. As shown in Fig. 5a, wild-type PP271 efficiently synthesized DNA primers of up to ∼12 nt when initiated with *ADP*. However, elongation of *ADP*–dC dimers was markedly inefficient in the presence of *NTPs*. Both Tyr^121^ variants displayed reduced primer synthesis with dNTPs compared to the wild-type and were more severely impaired when *NTPs* were used as substrates. These results indicates that Tyr^121^ does not simply acts as a sugar selector but also contribute to additional stabilization of the incoming nucleotide.

**Fig. 5.**
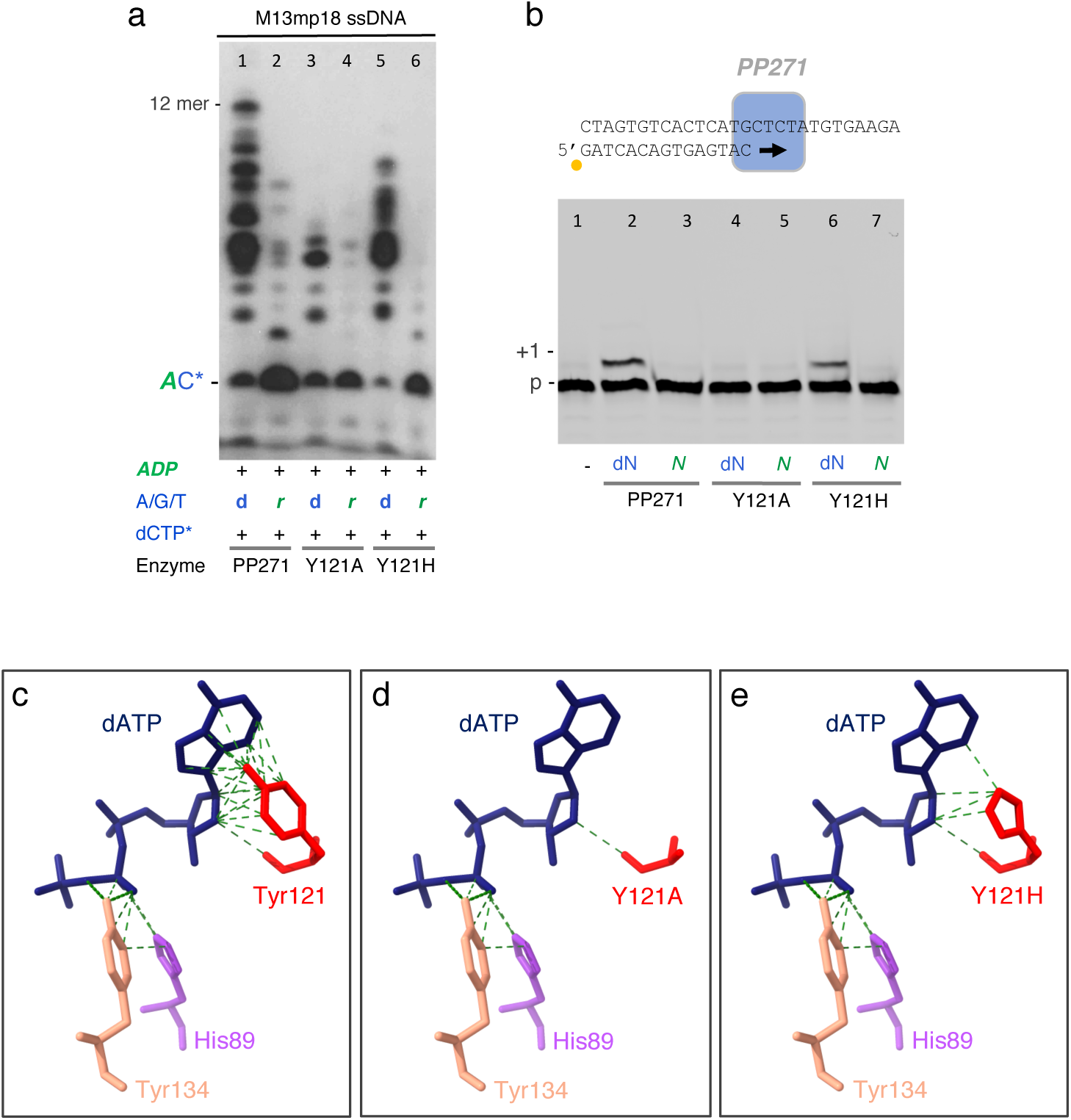
Tyr^121^ is required for nucleotide binding at the 3’site of PP271. (a) Primer synthesis by wild-type PP271 and the Y121A and Y121H variants (500 mM) in the presence of dNTP or *NTP* (100 µM each), [α-³²P]dCTP (16 nM) and *ADP* (100 µM) as the initiating nucleotide. (b) DNA polymerization activity of wild-type PP271 and the Y121A and Y121H mutants (500 mM) using a Cy5 labeled primer/template substrate (20 nM) and either dNTPs or *NTPs* (100 µM each). (c) AlphaFold model of PP271 showing the predicted interactions of Tyr^121^, Tyr^134^ and His^89^ (motif B) with dATP bound at the 3’nucleotide binding site. Tyr^121^ establishes multiple contacts with the adenine base and C1’ atom of the sugar moiety through its side chain and with C2’ atom of the sugar moiety through the backbone. (d) AlphaFold model of the Y121A variant showing the loss of most interactions with dATP. (e) AlphaFold modeled Y121H variant showing disruption of several interactions with dATP. In both variants, the backbone interaction with the C2′ atom of the sugar moiety is preserved.

Priming is a multistep process encompassing template recognition, dinucleotide formation, and subsequent extension that is limited under *in vitro* conditions that use labeled nucleotides. Thus, to specifically assess the role of Tyr^121^ in nucleotide incorporation during elongation, a DNA polymerase assay was performed, allowing direct evaluation of dNTP and *NTP* insertion independently of primer initiation. Using as DNA substrate a 5’-labeled primer annealed to a template, both wild-type PP271 and the Y121H mutant were capable of performing a +1 dNTP incorporation, whereas the Y121A mutant failed to do so (Fig. 5b). When *NTPs* were supplied, none of the three enzymes supported elongation. These results indicate that Tyr^121^ does not act as a strict steric gate preventing *NTP* incorporation. Instead, Tyr^121^ appears to contribute to stabilize the incoming nucleotide within the active site, thereby promoting efficient binding and catalysis. Consistent with this interpretation, a change of Tyr^121^ to histidine produces a milder defect, retaining partial activity.

Structural modeling using AlphaFold provided further insights into the role of Tyr^121^ and Tyr^134^ (Fig. 5c). Tyr^134^ is positioned in contact with the β-phosphate of the incoming nucleotide, suggesting a role unrelated to sugar discrimination. In contrast, Tyr^121^ is predicted to establish multiple interactions with the incoming nucleotide, including contacts with both the sugar and the nitrogenous base. Substitution of Tyr^121^ with histidine significantly reduces these interactions, while substitution with alanine leaves only minimal contact, primarily with the C2′ position of the sugar. These structural predictions are consistent with the experimental data and support a critical role for Tyr^121^ in nucleotide binding at the catalytic site, beyond a simple function as a sugar selector.

Remarkably, PP271 displayed very limited polymerase activity in the presence of Mg²⁺ as metal activator, extending the primer by just a single nucleotide. This observation indicates that the primase–polymerase balance of PP271 is strongly shifted toward primase activity, and likely correlates with its inability to generate primers longer than 12 nt. Together, these features highlight PP271 as an enzyme optimized for efficient primer generation rather than elongation, making it an attractive candidate for primer-free DNA amplification systems, prompting us to explore its functional interplay with the highly processive Phi29 DNApol.

### PP271-generated primers can be extended by Phi29 DNApol

The patented primer-free isothermal amplification system TruePrime (Picher et al., 2016) relies on DNA primer generation by *Tth*PrimPol, which are subsequently extended by Phi29 DNApol. In contrast to other PrimPols, and regardless of their preference for dNTPs, PP271 functions as a canonical primase, displaying limited DNA polymerase activity and utilizing Mg²⁺ as a metal cofactor, being thus compatible with Phi29 DNApol. Based on these properties, PP271 was considered a potential alternative for primer-free isothermal amplification systems.

To evaluate its suitability, the compatibility of PP271 with the high-fidelity Phi29 DNApol was assessed (Scheme, Fig. 6a). A pulse–chase experiment was designed to determine whether DNA primers synthesized by PP271 on an M13mp18 ssDNA template (pulse) could subsequently be extended by Phi29 DNApol (chase). The effect of adding *ADP* was tested to assess whether enhanced primase activity improves amplification efficiency and to evaluate any potential inhibitory effects on Phi29 DNApol.

**Fig. 6.**
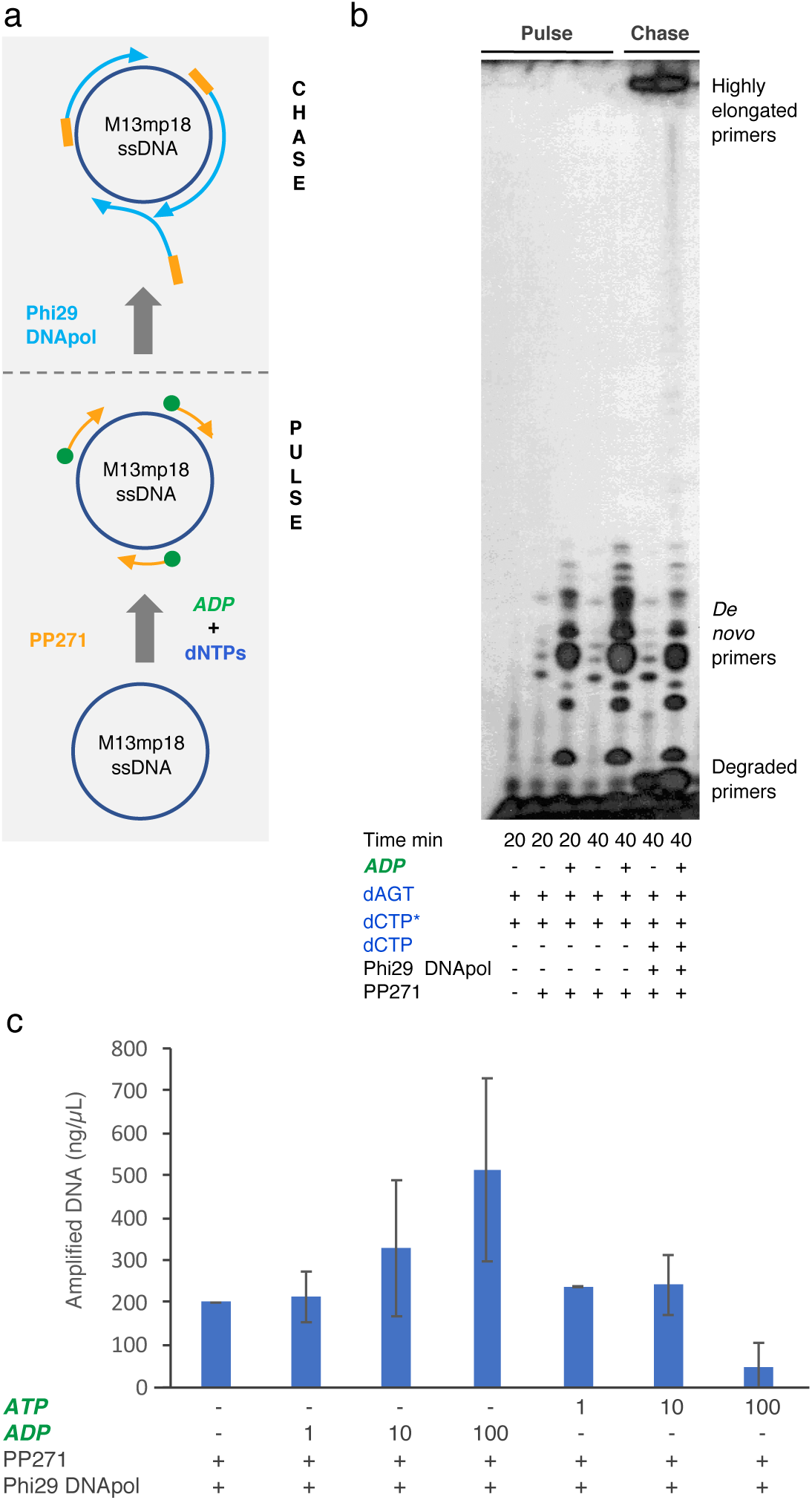
PP271 and Phi29 DNApol cooperate in hyperbranched rolling-circle amplification. (a) Schematic representation of the pulse-chase reaction shown in panel b. M13mp18 ssDNA was incubated with PP271 (500 nM) and dATP, dGTP, dTTP (100 µM each) and [α-³²P]dCTP (16 nM) and *ADP* (100 µM) as the initiating nucleotide, when indicated. Following a 20 min incubation at 30 °C to allow primer synthesis, Phi29 DNApol was added and the reaction was further incubated for 20 min. (b) Analysis of pulse-chase products. DNA primers synthetized during the pulse phase are detected at the bottom of the gel, whereas primers elongated by Phi29 DNApol migrates as high-molecular weight species near the top of the gel. (c) Quantification of primer-free amplification reactions performed with PP271 and Phi29 DNApol. M13mp18 ssDNA (1 ng) was amplified for 1 h at 30 °C in the presence of 1 mM dNTPs supplemented, where indicated, with 1, 10, or 100 µM *ADP* or *ATP*. Data represent the mean ± standard deviation from at least three independent experiments.

During the pulse phase, PP271 was allowed to synthesize primers on M13mp18 ssDNA in the presence of dNTPs, including radiolabeled dCTP, for 20 min at 30 °C (Fig. 6b, lanes 2 and 3). Phi29 DNApol was then added, and reactions were incubated for an additional 20 min (Fig. 6b, lanes 6 and 7). Control reactions lacking Phi29 DNApol were incubated for the same additional period (Fig. 6b, lanes 4 and 5).

As observed (Fig. 6b, lanes 2 *vs* 3; 4 *vs* 5; and 6 *vs* 7), the presence of *ADP* increased primer synthesis, consistent with previous results, generating products of approximately 12 nt in length. Phi29 DNApol was able to extend a fraction of these DNA primers, as evidenced by the appearance of high-molecular-weight products at the top of the gel. However, degradation products were also detected, likely due to the strong 3′→5′ exonuclease activity of Phi29 DNApol, as indicated by bands smaller than the PP271-synthesized primers. These results suggest that primers below a critical length (likely <6 nt) are inefficient substrates for elongation and are instead degraded, consistent with the known proofreading activity of Phi29 DNApol (Blanco et al., 1989). In this context, PP271-generated primers (∼10–15 nt) fall within the effective length range required for productive extension.

### Synergistic action of PP271 DNA primase and Phi29 DNApol enables efficient isothermal DNA amplification

Following confirmation of the synergistic activity between both enzymes, a primer-free DNA amplification assay was established. PP271 primase and Phi29 DNApol were evaluated in the presence or absence of increasing concentrations of *ADP* or *ATP* (Fig. 6c). Reactions containing low or no ribonucleotide supplementation produced comparable amplification yields of approximately 200 ng/µl, indicating that basal activity can be achieved under minimal conditions.

Remarkably, higher *ADP* concentrations significantly enhanced amplification efficiency, reaching up to ∼600 ng/µl at 100 µM *ADP*. In contrast, increasing *ATP* concentrations led to a progressive decline in amplification output. A plausible explanation is that *ATP* may be misincorporated at the elongation site of Phi29 DNApol, thereby interfering with its catalytic cycle. Exonucleolytic removal of the incorrectly incorporated *ATP* would be required to allow a correct dNTP addition, reducing the overall efficiency of DNA synthesis.

Conversely, *ADP* appears to selectively stimulate the primase activity of PP271 without compromising the functionality of Phi29 DNApol. This dual effect positions *ADP* as a favorable cofactor for this amplification system, enhancing primer synthesis while maintaining efficient DNA elongation. Collectively, these findings highlight the potential of the PP271–Phi29 DNApol enzymatic pair as a robust platform for primer-independent isothermal DNA amplification.

## Discussion

PrimPols are members of the archaeo-eukaryotic primase (AEP) superfamily characterized by their dual primase–polymerase activity, which enables both *de novo* primer synthesis and primer extension. These enzymes are widely distributed across all domains of life, including archaea, eukaryotes, bacteria, and viruses (Blanco et al., 2019). In bacteria, however, PrimPol genes are frequently associated with mobile genetic elements, suggesting extensive horizontal gene transfer during evolution (Sanchez-Berrondo et al., 2012)(García-Quintans et al., 2020)

A unique feature of PrimPols is their ability to synthesize DNA primers; however, they may vary in the nature of the sugar of the primer’s initiating nucleotide. Moreover, they exhibit substantial differences in their DNA polymerization capabilities. The PrimPol from the archaeal plasmid pRN1 (firstly named PrimPol) and that from *Thermus thermophilus* display highly processive polymerase activities and are capable of synthesizing DNA tracts of several kilobases (Lipps, 2004; Picher et al., 2016). Likewise, the PrimPols encoded by the Deep-sea phage NrS-1 and Mimivirus function as *bona fide* replicative enzymes, providing these viruses with an increased degree of independence from their host replication machinery (Gupta et al., 2019; Zhu et al., 2017). In contrast, human PrimPol exhibits a much more limited polymerase activity, although it can still elongate primers by several hundred nucleotides under appropriate conditions (García-Gómez et al., 2013). Notably, genetic studies have demonstrated that PrimPoĺs primase activity, rather than its polymerase activity, is the essential function required *in vivo* for replication stress tolerance and genome maintenance (Mourón et al., 2013; Torregrosa-Muñumer et al., 2017).

The biochemical properties of PP271 further widens this functional spectrum. Our results show that PP271 possesses a very limited DNA polymerase activity, being only capable of extending a pre-existing primer by a single nucleotide in the presence of Mg²⁺ (Fig. 5b). Moreover, its primase activity generates exceptionally short primers, typically not exceeding 12 nt (Fig. 4a). This behavior contrasts sharply with previously characterized PrimPols that efficiently combine primer synthesis with extensive DNA elongation.

Interestingly, the EFAS271 genome encodes not only PP271 but also a canonical DnaG-like primase and a family B DNA polymerase. This genetic organization suggests a division of labor during viral DNA replication. DnaG-synthesized *RNA* primers could be responsible for initiating genome replication at a specific origin sequence, potentially in association with the phage-encoded helicase, whereas the family B DNA polymerase would perform processive genome synthesis. Within this framework, PP271 may fulfill a specialized accessory function rather than serving as an essential replicative enzyme. One possibility is that PP271 participates in DNA repair pathways, helping the phage counteract bacterial defense systems that damage viral DNA. Alternatively, it may play a canonical PrimPol-like role, facilitating replication restart, thus allowing damage tolerance. Such a role would be particularly advantageous considering the remarkably short biosynthetic phase of the EFAS271 life cycle, where virion assembly is completed within approximately eight minutes (Topka-Bielecka et al., 2020).

Despite their functional diversity, PrimPols share a conserved catalytic AEP core containing the canonical motifs A, B, and C involved in the two metal-ion catalysis mechanism (Iyer et al., 2005). In contrast, their C-terminal regions are highly variable and have evolved distinct structural modules involved in primer initiation. Examples include the Zn-finger domain of human (García-Gómez et al., 2013; Iyer et al., 2005) and murine PrimPol (Carvalho et al., 2025); the helix bundle domain (HBD) of the pRN1 PrimPol (Beck et al., 2010; Boudet et al., 2019), and the PriCT domain present in *Tth*PrimPol (Picher et al., 2016) and PP271. These accessory domains play a crucial role during primer synthesis by interacting with the initiating nucleotide that eventually becomes the 5′ end of the nascent primer. Functionally, they resemble the C-terminal domain of the large subunit of eukaryotic heterodimeric primases, which stabilizes the initiating nucleotide during primer formation (Baranovskiy et al., 2016).

In human PrimPol, the Zn-finger domain interacts directly with the initiating nucleotide through contacts with its phosphate moiety, particularly the γ-phosphate (Martínez-Jiménez et al., 2018). More recently, Boldinova et al. (2023) proposed that Arg^417^ of human PrimPol may perform a role analogous to that of Arg^306^ in the p58C subunit of the eukaryotic Polα–primase complex (Boldinova et al., 2023). Current models suggest that this interaction between the accessory domain and the 5′ terminal nucleotide is transient and persists only until the primer reaches a sufficient length for productive hand-off to the replicative polymerase. This process likely involves conformational rearrangements that separate the catalytic AEP core from the accessory domain through a flexible linker region.

Structural modeling of the PP271 PriCT domain using AlphaFold (Fig. 7b) revealed important differences relative to previously characterized initiation domains. The electrostatic surface of the putative initiating nucleotide-binding pocket appears substantially less positively charged than that observed in the p58C domain (Fig. 7a). In the latter, the α- and γ-phosphates of *ATP* are stabilized by a network of hydrogen bonds involving Arg^302^, His^303^ and Arg^306^ (Baranovskiy et al., 2016). These residues create a positively charged pocket that efficiently accommodates the negatively charged triphosphate group. In contrast, the corresponding region in PP271 contains Arg^186^, Asn^187^, and two glutamate residues (Glu^183^ and Glu^227^). The presence of two negatively charged residues within the putative phosphate-binding pocket immediately suggested that *ATP* binding might be energetically unfavorable due to electrostatic repulsion between the glutamate side chains and the triphosphate moiety. This structural feature provides a plausible explanation for the unexpected preference of PP271 for diphosphate nucleotides as the initiating nucleotide. Whereas *ATP* may be insufficiently stabilized within the pocket (Fig. 7c), *AMP* would likely lack the phosphate interactions required to support productive initiation. *ADP* may therefore represent an optimal compromise that enables stabilizing interactions without imposing excessive electrostatic constraints (Fig. 7d). Such a mechanism could explain both the nucleotide preference observed experimentally and the unusually short primers synthesized by PP271. The weak interaction predicted between PriCT and the 5′ nucleotide may further suggest that PP271 is not optimized for maintaining prolonged contact with growing primers. Consequently, the enzyme may not display the strong 5′-end stabilization characteristic of PrimPols that synthesize long DNA products, such as human PrimPol or the *T. thermophilus* ortholog. Alternatively, the *ADP*-initiated short DNA primers preferentially made by PP271 may be an indirect consequence of its inability to efficiently engage the γ-phosphate of the initiating nucleotide, implying a reduced primer stabilization during elongation.

**Fig. 7.**
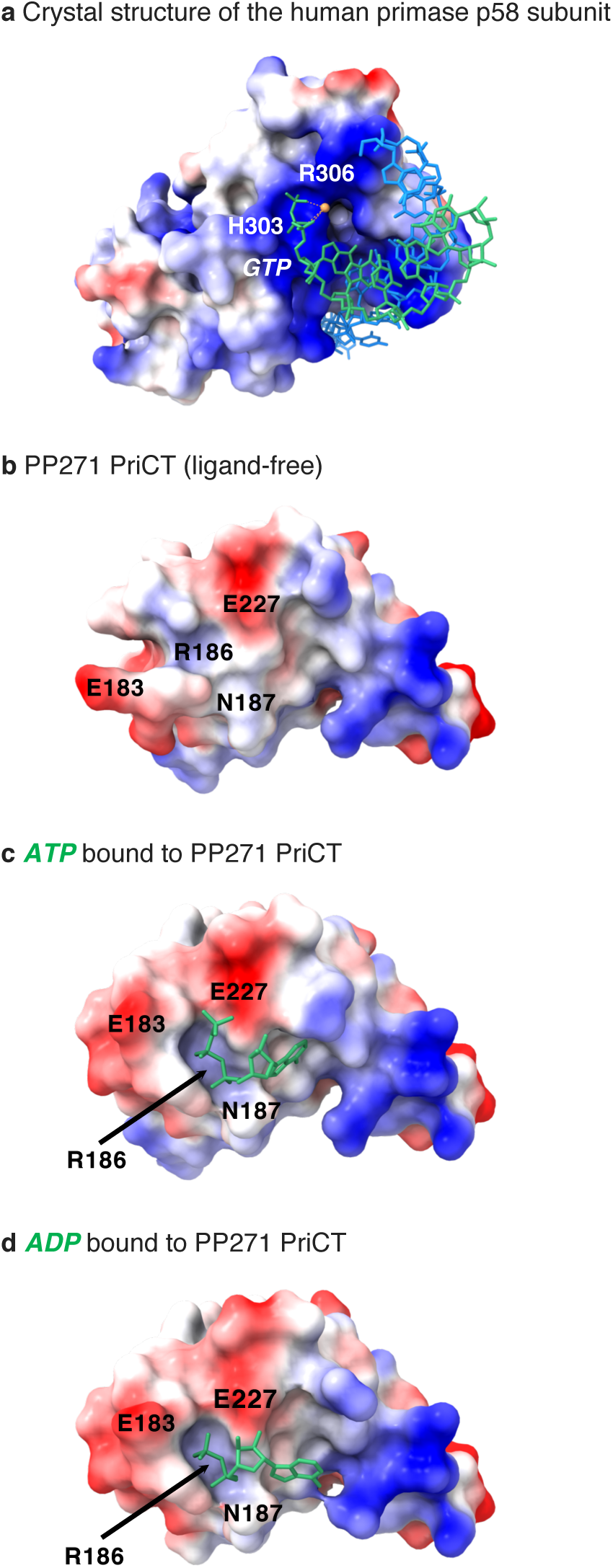
(a) Crystal structure of the C-terminal domain of the human DNA primase large subunit (p58) in complex with a DNA template–RNA primer duplex (PDB: 5F0Q). (b) Surface representation of the PP271 PriCT domain predicted by AlphaFold3. (c and d) AlphaFold3-predicted models of *ATP* (c) and *ADP* (d) bound to the PP271 PriCT domain.

The pronounced imbalance between the primase and polymerase activities of PP271 creates attractive opportunities for biotechnological applications. Because its contribution to DNA synthesis in a primer-less amplification method is largely restricted to the generation of short DNA primers, PP271 can be exploited to minimize the incorporation of synthesis errors associated with extensive DNA polymerization. In fact, PP271 was shown to be a good alternative to *Tth*PrimPol for SARS-CoV2 *RNA* detection through padlock-probe amplification in combination with a novel Phi29 DNApol variant (Blanco, Luis et al., n.d.; Prado et al., 2025).

In the present work, we further demonstrate that primers synthesized by PP271 can efficiently support amplification of M13mp18 ssDNA through hyperbranched rolling-circle amplification (HRCA). The observed synergy between PP271 and Phi29 DNApol expands the toolbox of isothermal amplification methods and may offer an additional advantage in terms of fidelity. Since PP271 contributes only short initiation products and the bulk of DNA synthesis is performed by the highly accurate Phi29 DNApol, the overall amplification process may achieve a higher fidelity than systems employing more processive PrimPols, such as the *Tth*PrimPol that is being used in the TruePrime™ DNA amplification system (Picher et al., 2016).

## Author contribution

LB and MIMJ did the conceptualization of the study. AJM, SGG, RG, IL and MIMJ conducted the experiments. AJM and MIMJ performed data analysis and wrote original draft of the manuscript. The final version was reviewed by AJM, MIMJ and LB. The resources and the funding acquisition were provided by LB. All authors read and approved the final version of the manuscript.

## Statements and Declarations

### Ethics approval

This article does not contain any studies with human participants or animals performed by any of the authors.

